# A novel Vps34 complex constrains loop extrusion-mediated P-H2A spreading

**DOI:** 10.1101/2022.03.09.482075

**Authors:** Hui Chen, Tingting Feng, Simu Liu, Wei-Guo Zhu

## Abstract

The establishment of appropriate phosphorylated histone H2AX (γH2AX) foci is required for the precise initiation of DNA damage response (DDR) signaling, which contributes to the efficient repair of DNA double-strand breaks (DSB)^1^. Therefore, it is important to understand exactly how γH2AX foci are established in response to DSB. Recently, an attractive mechanism has been proposed to explain this process^2^. DSB-anchored cohesin-mediated loop extrusion promotes H2AX-containing chromatin to pass through DSB-recruited ATM, by which H2AX is phosphorylated to form γH2AX foci on each side of the DSB. The loop extrusion-promoted enlargement of γH2AX-modified chromatin is halted upon cohesin encountering boundary factors such as CTCF. This cohesin-dependent mechanism was shown to be conserved among eukaryotic cells, because deletion of the cohesin regulator Pds5 in budding yeast also results in extended spreading of phosphorylated H2A at Ser129 (P-H2A)^2^. However, yeast lacks CTCF or CTCF-like boundary factors that restrict cohesion-mediated loop extrusion. Here, we report a novel vacuolar sorting protein 34 (Vps34) complex, which might function as a boundary factor that constrains cohesin-dependent loop extrusion-mediated P-H2A spreading.

## RESULTS

To explore the roles of yeast Vps family proteins in the DDR, we performed a targeted genetic screening. In total, 41 *VPS* gene deletion strains were screened in the presence of the radiomimetic compound zeocin, which induces DSB of these strains; of these strains, 18 *vps* mutants displayed greater sensitivity than the wild type to zeocin. Among them, *vps34Δ* and *vps15Δ* interested us greatly, not only because they showed hypersensitivity to zeocin (Figure S1A) but also because Vps34 is the sole phosphatidylinositol-3-kinase in yeast and plants and is widely conserved, from yeast to humans^3^. Vps34 phosphorylates phosphatidylinositol (PI) at the 3-hydoxyl group of the inositol ring to generate PI(3)P^4^, and the putative serine/threonine kinase Vps15 is required for its kinase activity^3^. In view of the fact that PI(3)P can be further phosphorylated by Fab1 to form PI(3, 5)P_2_^5^, we observed the phenotypes of *fab1Δ* and the Fab1 regulator *vac7Δ* and *vac14Δ* mutants in the presence of zeocin, and we found that the deletion of these genes did not affect the sensitivity of yeast to zeocin (Figure S1B). Furthermore, abolishment of the kinase activity of Vps34 or Vps15 was unable to complement their corresponding mutation phenotypes (Figure S1C, D). These results demonstrate a key role for PI(3)P or Vps34 kinase activity in the regulation of the DDR.

In budding yeast, the Vps34-Vps15 kinase module is known to function in autophagy and vacuolar protein sorting when associated with Vps30-Atg14 and Vps30-Vps38 to form complexes I and II, respectively^3^. Unlike Vps34 and Vps15, a lack of Vps30, Vps38, or Atg14 protein did not alter the sensitivity of yeast to zeocin (Figure S1E), indicating that these two known protein complexes are not involved in the DDR. In consideration of the key role Vps34-associated proteins play in facilitating Vps34’s accomplishment of its cellular functions, it is reasonable to identify a previously unknown Vps34 complex involved in the DDR.

To investigate the effects of Vps34 or Vps15 deficiency on the DDR, we investigated the levels of P-H2A in wild type and mutant cells. Neither Vps34 deletion nor Vps15 deletion in the absence of zeocin significantly altered P-H2A levels. Nevertheless, a marked increase of zeocin-induced P-H2A in both *vps34Δ* and *vps15Δ* mutants was observed in comparison with P-H2A in the wild type. Consistent with the key role played by Vps34 in generating PI(3)P, more P-H2A was detected in *vps34Δ* mutants than in *vps15Δ* mutants (Figure 1A).

**Figure 1.**
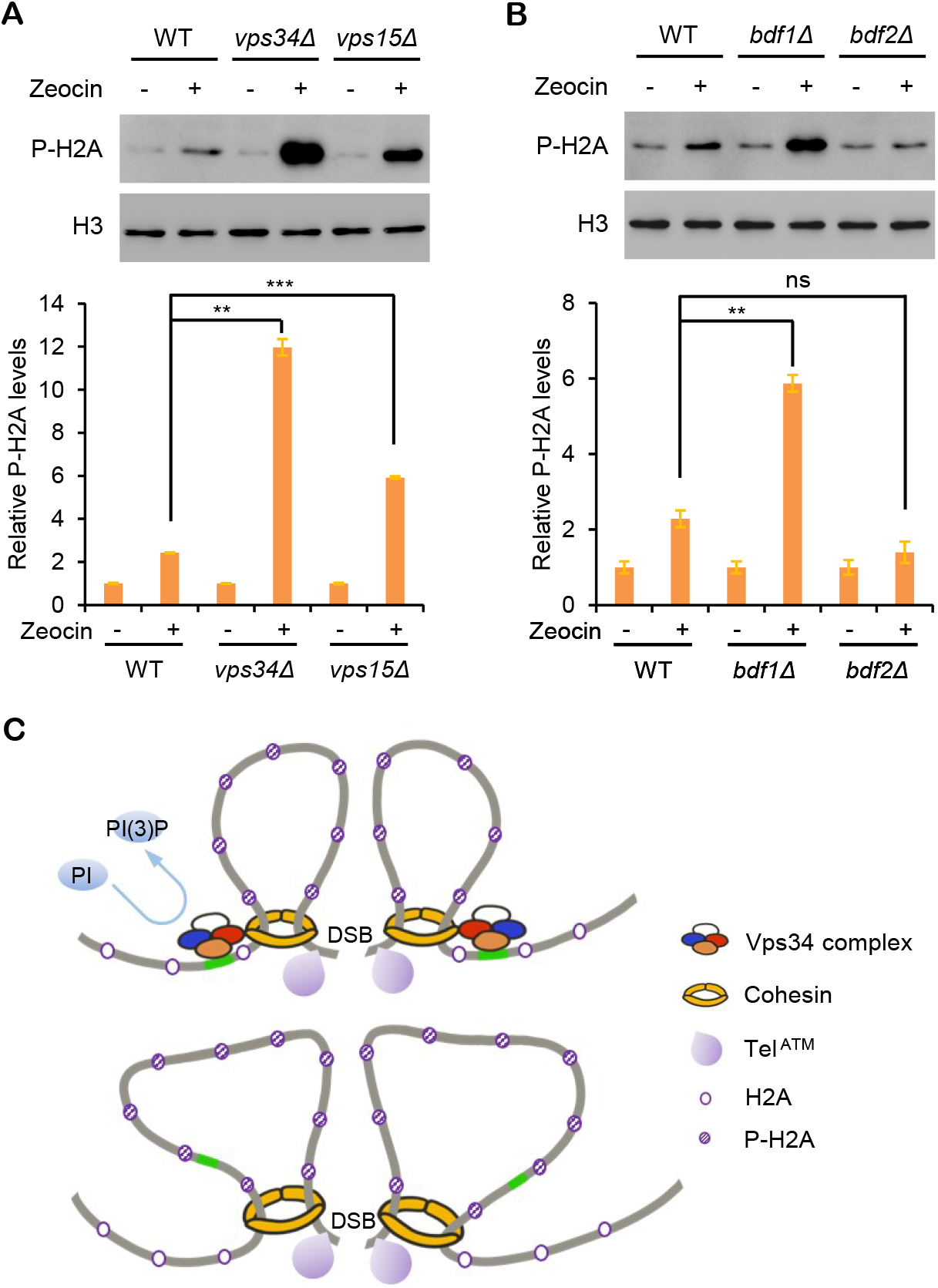
The Vps34 complex functions as a boundary factor that constrains DSB-triggered H2A phosphorylation. (A,B) Yeast cells, comprising *vps34Δ, vps15Δ* mutants (A) or *bdf1Δ, bdf2Δ* mutants (B) and wild type (WT) cells were treated with or without zeocin for 1 hour. Protein samples from the indicated strains were analyzed using anti-P-H2A antibodies. H3 was used as a loading control. Three biological repeats were performed and analyzed. \*\**p* < 0.01; \*\*\**p* < 0.001; NS, no statistically significant difference. Data are shown as mean ± standard deviation (SD). (C) A hypothetical working model for the Vps34 complex (Vps34-Vps15-Bdf1-Pds5 protein complex) in restricting DSB-triggered P-H2A spreading. In strains with the Vps34 complex (upper panel), cohesin-mediated loop extrusion could be halted normally, thus phosphorylation of H2A is limited. In strains without the Vps34 complex (lower panel), cohesin-mediated loop extrusion could not be halted efficiently, thus more H2A is phosphorylated.

These results reminded us of mammalian Brd4, the knockdown of which also causes an increase in ionizing-radiation-induced γH2AX^6^. Similar to Brd4, which is a bromodomain and extra-terminal (BET) family member, both Bdf1 and Bdf2 in yeast contain the BET domain. Interestingly, the *Bdf1Δ* but not the *Bdf2Δ* mutant displayed more sensitivity than the wild type in response to zeocin (Figure S1F), while the level of P-H2A induced by zeocin in the Bdf1-deficient strain increased remarkably compared with that seen in the wild type and was comparable to that of the Vps15-deficient mutant (Figure 1B). Given the similar phenotypes of *vps34Δ*, *vps15Δ*, and *Bdf1Δ* in response to zeocin, and the interaction of Vps34 with additional proteins for its cellular functions, we speculated that Bdf1 might form a protein complex with Vps34 and Vps15. To test this possibility, we constructed yeast strains with tagged proteins and performed co-immunoprecipitation assays. Bdf1 could be co-precipitated with both Vps34 and Vps15 proteins (Figure S2A, B), suggesting they are in the same protein complex that is involved with restricting P-H2A accumulation.

We next asked how the Vps34-Vps15-Bdf1 complex modulates P-H2A. Previous studies have reported that Brd4 functions as an endogenous insulator that constrains ionizing-radiation-induced increase of γH2AX by recruiting the condensin complex to remodel chromatin^6^. In yeast, the cohesin regulator Pds5 has also been shown to modulate DSB-triggered P-H2A establishment^2^. We next analyzed whether Vps34 or Bdf1 could associate with condensin complex or Pds5. Intriguingly, both Vps34 and Bdf1 could associate with Pds5 in our assays (Figure S2C, D). Consistent with the role played by Pds5 in the stabilization of cohesin binding to chromosomes^7^, the deletion of either Vps34 or Bdf1 led to the noticeable removal of cohesin from chromatin (Figure S2E). These results revealed that Vps34, Vps15, Bdf1, and Pds5 are likely to form a previously unknown Vps34 complex for the regulation of cohesin-mediated P-H2A establishment.

## SUMMARY

In mammalian cells, Pds5 is required to fulfill the boundary function of CTCF, with Pds5 deficiency resulting in longer loops^8^. Furthermore, cohesin-mediated loop extrusion can be constrained even after the deletion of CTCF^9^, implying the existence of additional boundaries other than just CTCF-bound loci. In yeast cells, Pds5 also limits the size of cohesin-mediated loops, even though no CTCF orthologue is known to exist in yeast^10^, indicating a conserved boundary role of Pds5 in loop extrusion. Taking our findings together, we propose that the Pds5-including Vps34 complex we have discovered might function as a boundary for cohesin-dependent loop extrusion and thus act to constrain P-H2A spreading (Figure 1C). Future studies may provide more evidences which are involved in the regulation of cohesin-mediated loop extrusion, for example, by comparison of the chromatin loop patterns between *vps34Δ* and wild type yeast using bioinformatic methods such as Hi-C (high through chromosome conformation capture).

## ACKNOWLEDGMENTS

This work was supported by the Human Resources and Social Security Administration of Shenzhen Municipality (to H.C.), the National Key R&D program of China (2017YFA0503900), the National Natural Science Foundation of China (81720108027), the Science and Technology Innovation Commission of Shenzhen Municipality (JCYJ20200109114214463), and the Shenzhen Bay Laboratory (SZBL2019062801011). We thank Prof. Junbiao Dai (Shenzhen Institutes of Advanced Technology, Chinese Academy of Sciences) for providing the BY4741 strain, Xi Zhao (Institute of Microbiology, Chinese Academy of Sciences) for discussing the methods, and Dr. Dachuan Lin (Shenzhen University) for sharing the cell disruptor instrument.

## AUTHOR CONTRIBUTIONS

Conceptualization: H.C.; Methodology: H. C.; Investigation: H.C. and T.T.F.; Data Analysis and Discussion: H.C., S.M.L. and W.G.Z.; Writing-Original Draft: H.C.; Writing-Review and Editing: H.C. and W.G.Z. Funding Acquisition: W.G.Z.

