## Supplemental Information for "A novel Vps34 complex constrains loop extrusion-mediated P-H2A spreading"

Figure S1. Phenotypes of the indicated strains in response to zeocin.

Figure S2. Identification of a novel Vps34 complex.

Supplemental Experimental Procedures

**Figure S1. Phenotypes of the indicated strains in response to zeocin.**

(A-F) Ten-fold serial dilutions of the indicated strains were spotted onto YPD medium with or without zeocin. (A) Deletion of Vps34 or Vps15 both resulted in hypersensitivity to zeocin. (B) Deficiency of Fab1 and its regulators in yeast cells did not alter their sensitivity to zecocin. (C) Destruction of the kinase activity of Vps34 could not rescue *vps34Δ* phenotypes. (D) Destruction of the kinase activity of Vps15 could not rescue *vps15Δ* phenotypes. (E) Deficiency of proteins that form known Vps34 complexes did not alter their sensitivity to zeocin. (F) Deletion of Bdf1 but not Bdf2 resulted in greater sensitivity to zeocin than that seen in the wild type.

**Figure S2. Identification of a novel Vps34 complex.**

(A-B) Coimmunoprecipitation of Bdf1 with Vps34 (A) or Vps15 (B). Anti-GFP antibodies were used to immunoprecipitate Vps34-GFP or Vps15-GFP, and coimmunoprecipitation of Bdf1-mCherry was analyzed by immunoblotting. IgG was used as a control for immunoprecipitation.

(C) Coimmunoprecipitation of Pds5 with Vps34. Vps34-GFP was immunoprecipitated, and coimmunoprecipitation of Pds5-mCherry was analyzed by immunoblotting.

(D) Coimmunoprecipitation of Bdf1 with Pds5. Pds5-GFP was immunoprecipitated, and coimmunoprecipitation of Bdf1-mCherry was analyzed by immunoblotting.

(E) Immunoblot analysis of the cohesin subunit Mcd1 in the whole cell extract and the chromatin pellet fraction from the indicated strains. Anti-GFP antibodies were used to detect Mcd1-GFP. GAPDH and H3 were used as loading controls.

### Supplemental Experimental Procedures

*Media, growth conditions, and yeast strains*

Yeast cells were grown in solid yeast extract-peptone-dextrose (YPD) or synthetic drop-out (SD) medium for 3-5 days at 30ºC or were cultured in liquid medium overnight. All *Saccharomyces cerevisiae* strains used in this study were derivatives of BY4741 (*MATa his3Δ1 leu2Δ0 met15Δ0 ura3Δ0*) background. All deletion strains were from the Yeast Knockout MATa Haploid Collection (Transomic). Introduction of an epitope tag to indicate proteins was performed using a one-step PCR-based strategy[S1], and the yeast strains were verified by PCR testing, DNA sequencing, and immunoblot analysis.

*Plasmid construction and transformation*

Full-length Vps34 or Vps15 was amplified by PCR and introduced into the vector pGADT7 ($Leu^+$) (Clontech) using the homologous recombination method to generate an expression plasmid. Substitution mutations were generated by PCR-mediated site-directed mutagenesis. All constructs were confirmed by DNA sequencing. Yeast transformation was performed according to the protocol in the Yeastmaker™ Yeast Transformation System 2 user manual (Clontech).

*DNA damage sensitivity assay*

Yeast strains were cultured on YPD plates for 3 days then diluted to $OD_{600}$ = 1.0 in sterile water with 0.9% (w/v) NaCl. Ten-fold serial dilutions were prepared and 2 µl volumes were spotted onto plates with or without 10 µg/ml zeocin. Images were taken after incubation for 3 days.

*Protein extraction and immunoblot analysis*

Protein was extracted as previously described with a few modifications[S2]. Cells were collected by centrifugation at 845 × g for 1 min, then washed once with 500 µl of 20% (w/v) trichloroacetic acid (TCA). Next, cells were resuspended in 200 µl of 20% TCA and lysed by disruption with 0.5-mm-diameter zirconia beads using a cell disruptor instrument (power = 64 Hz, time = 4 × 45s). The zirconia beads were washed twice with 200 µl of 5% TCA each time, and the protein pellets were collected by centrifugation at 845 × g for 10 min. The pellets were resuspended in 2×SDS-PAGE loading buffer and the extracts were then neutralized by adding 1 M Tris base and boiling for 10 min. The proteins were separated on SDS-PAGE gels and then immunoblotted with antibodies. The antibodies used in this study were: anti-H2AS129ph (ab15083, 1:1000); anti-H3 (ab1791, 1:1000); anti-GFP (MBL D153-3 and Roche 11814460001, 1:1000); anti-mCherry (ABB-A02080, 1:1000); anti-GAPDH (Proteintech 60004-1-lg, 1:5000); and IgG (Beyotime A7028).

*Immunoprecipitation*

Immunoprecipitation assays were performed as previously descripted, with a few modifications[S3]. The yeast cells were harvested during log-phase and washed twice with water. Cells were resuspended with 300 µl lysis buffer (50 mM HEPES-NaOH, pH 7.5, 150 mM NaCl, 1 mM EDTA, 1 mM DTT, 1 mM PMSF, 0.05% NP-40, 10% glycerol, 1 × Roche protease inhibitor cocktail) and were lysed by disruption with 0.5-mm-diameter zirconia beads using a cell disruptor instrument (power = 64 Hz, time = 4 × 45s). The cell lysates were cleaned by centrifugation at 13,500 × g at 4ºC for 10 min. The supernatants were incubated with GFP antibodies at 4ºC. After overnight incubation, protein G sepharose (GE Healthcare) was added and incubated for an additional 4 h, the beads were then washed five times with lysis buffer. The beads were resuspended in 1×SDS-PAGE loading buffer, and any proteins that were bound to the beads were eluted by boiling for 10 min, before being analyzed by western blot.

*Chromatin fractionation*

Yeast strains were grown to log-phase in liquid YPD medium. Chromatin was extracted as previously described, with a few modifications[S4]. Cells were harvested and washed twice with PBS buffer. Spheroplasts were obtained by zymolase digestion (50 mM KPO4 buffer pH7.4, 0.6 M sorbitol, 10 mM DTT, and 20 U zymolase-20T) for 30 min at 35ºC, then washed once with Spheroplast Washing Buffer (50 mM HEPES, pH 7.5, 150 mM NaCl, 0.4 M sorbitol, 1 mM PMSF, and 1 × Roche protease inhibitor cocktail). Spheroplasts were resuspended in 500µl Lysis Buffer A (50 mM HEPES, pH 7.5, 150 mM NaCl, 1 mM DTT, 1 mM ATP, 1 mM PMSF, 1 × Roche protease inhibitor cocktail, and 0.25% triton X-100) for 10 min on ice to obtain whole cell lysate, then the whole cell lysate was slowly laid onto 500µl Lysis Buffer A-S (50 mM HEPES, pH 7.5, 150 mM NaCl, 1 mM DTT, 1 mM ATP, 1 mM PMSF, 1 × Roche protease inhibitor cocktail, 0.25% triton X-100, and 30% sucrose), and centrifuged at 13,500 × g at 4ºC for 10 min. The chromatin pellets were obtained and washed with Chromatin Washing Buffer (50 mM HEPES, pH 7.5, 150 mM NaCl, 1 mM DTT, 1 mM ATP, 1 mM PMSF, and 1 × Roche protease inhibitor cocktail). Proteins that bound to the chromatin were eluted by ultrasonic processing (power, 20%; 2 sec on, 4 sec off, 10 cycles) in SDS-PAGE loading buffer, and subjected to immunoblot analysis.

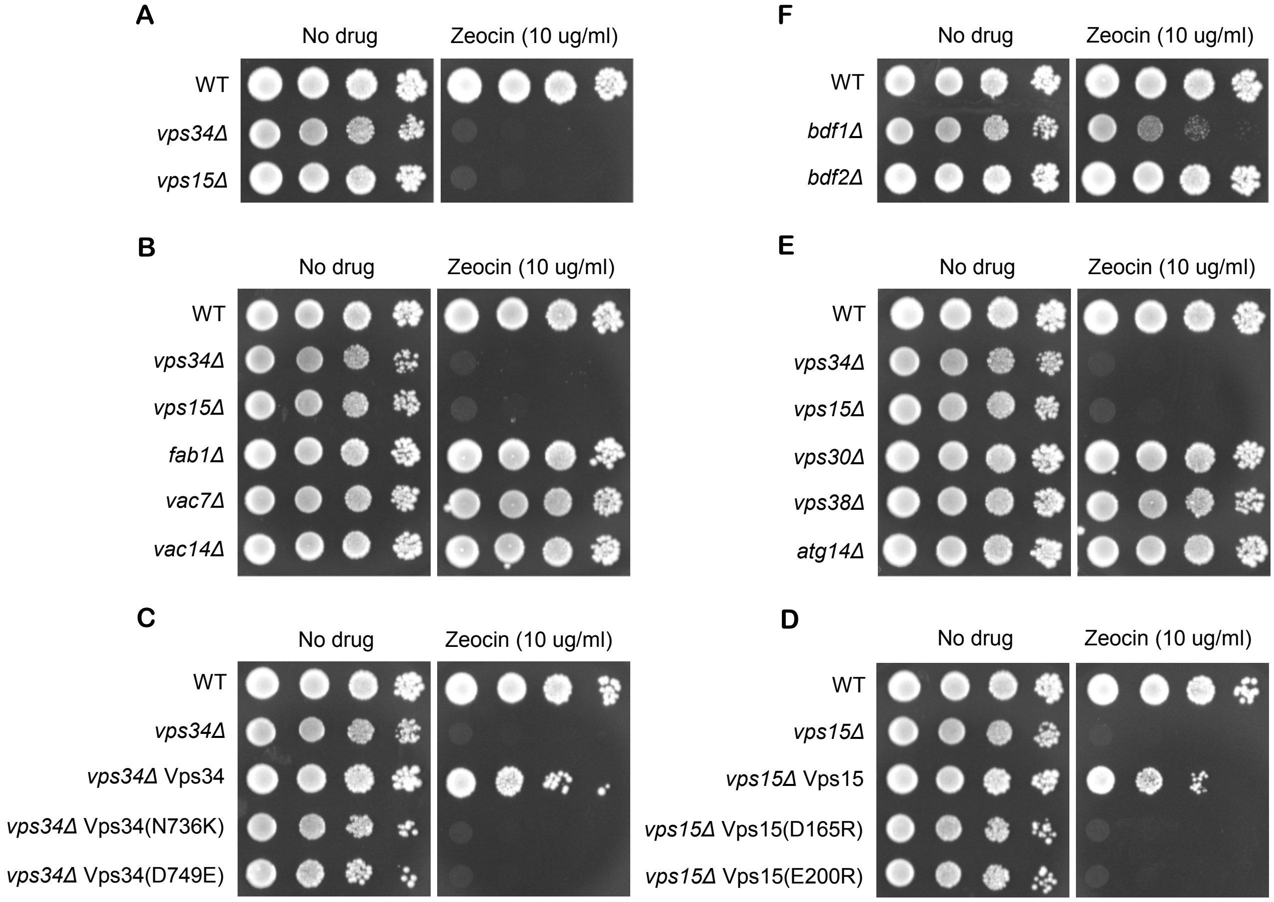


**Figure S1. Phenotypes of the indicated strains in response to zeocin.**
(A-F) Ten-fold serial dilutions of the indicated strains were spotted onto YPD medium with or without zeocin. (A) Deletion of Vps34 or Vps15 both resulted in hypersensitivity to zeocin. (B) Deficiency of Fab1 and its regulators in yeast cells did not alter their sensitivity to zecocin. (C) Destruction of the kinase activity of Vps34 could not rescue *vps34Δ* phenotypes. (D) Destruction of the kinase activity of Vps15 could not rescue *vps15Δ* phenotypes. (E) Deficiency of proteins that form known Vps34 complexes did not alter their sensitivity to zeocin. (F) Deletion of Bdf1 but not Bdf2 resulted in greater sensitivity to zeocin than that seen in the wild type.

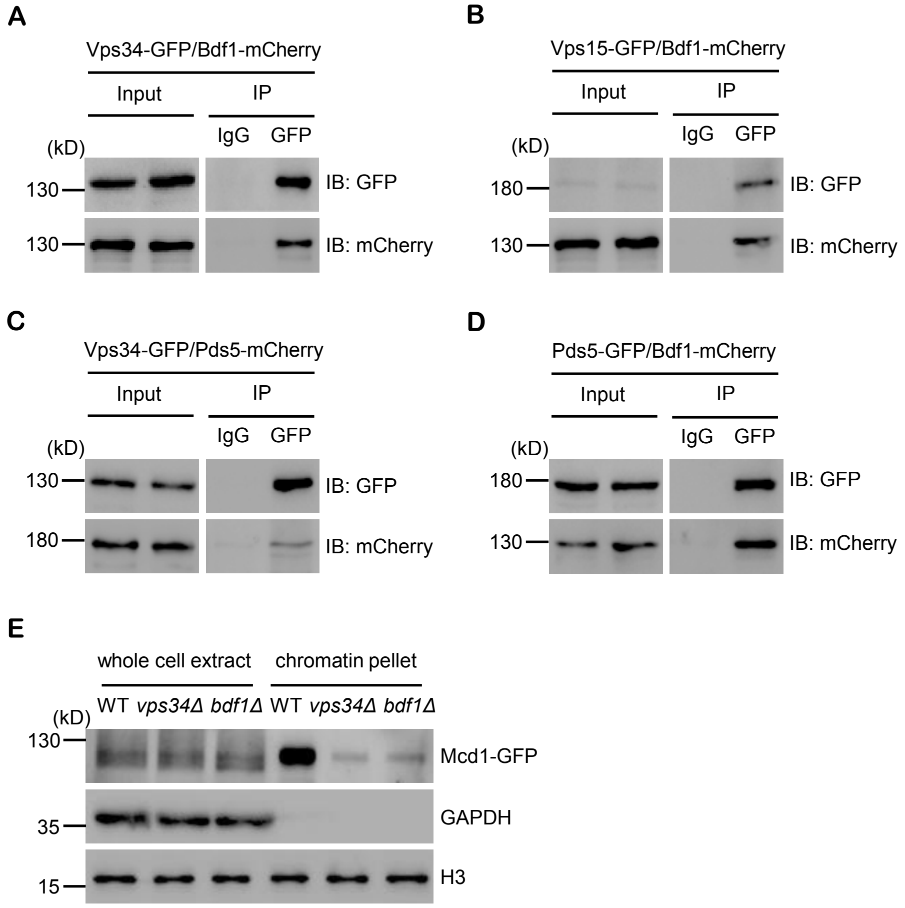


**Figure S2. Identification of a novel Vps34 complex.**
(A-B) Coimmunoprecipitation of Bdf1 with Vps34 (A) or Vps15 (B). Anti-GFP antibodies were used to immunoprecipitate Vps34-GFP or Vps15-GFP, and coimmunoprecipitation of Bdf1-mCherry was analyzed by immunoblotting. IgG used as a control for immunoprecipitation.
(C) Coimmunoprecipitation of Pds5 with Vps34. Vps34-GFP was immunoprecipitated, and coimmunoprecipitation of Pds5-mCherry was analyzed by immunoblotting.
(D) Coimmunoprecipitation of Bdf1 with Pds5. Pds5-GFP was immunoprecipitated, and coimmunoprecipitation of Bdf1-mCherry was analyzed by immunoblotting.
(E) Immunoblot analysis of the cohesin subunit Mcd1 in the whole cell extract and the chromatin pellet fraction from the indicated strains. Anti-GFP antibodies were used to detect Mcd1-GFP. GAPDH and H3 were used as loading controls.
